# Characterizing Spatially Continuous Variations in Tissue Microenvironment through Niche Trajectory Analysis

**DOI:** 10.1101/2024.04.23.590827

**Authors:** Wen Wang, Shiwei Zheng, Sujung Crystal Shin, Guo-Cheng Yuan

**Affiliations:** Department of Genetics and Genomic Sciences, Icahn School of Medicine at Mount Sinai, New York, NY, USA

## Abstract

Recent technological developments have made it possible to map the spatial organization of a tissue at the single-cell resolution. However, computational methods for analyzing spatially continuous variations in tissue microenvironment are still lacking. Here we present ONTraC as a strategy that constructs niche trajectories using a graph neural network-based modeling framework. Our benchmark analysis shows that ONTraC performs more favorably than existing methods for reconstructing spatial trajectories. Applications of ONTraC to public spatial transcriptomics datasets successfully recapitulated the underlying anatomical structure, and further enabled detection of tissue microenvironment-dependent changes in gene regulatory networks and cell-cell interaction activities during embryonic development. Taken together, ONTraC provides a useful and generally applicable tool for the systematic characterization of the structural and functional organization of tissue microenvironments.

## Main

Cellular behavior is determined not just by its intrinsic identity but also by its interactions with the external tissue microenvironment. Systematic analysis of tissue microenvironment variations can provide mechanistic insights into gene regulation during development and in diseases. Spatial transcriptomics offers an opportunity to dissect these variations. The prevailing approaches involve the identification of spatial domains as discrete organizational units^1–8^. The segmentation of tissue into spatial domains serves as a useful guide to study cell-state variations associated with distinct regions. However, a limitation is that these methods cannot capture the continuous tissue microenvironment variations.

Spatial trajectory analysis aims to model spatially continuous variations. Current methods for constructing spatial trajectories can be viewed as extensions of pseudotime analysis^9–13^, which were originally developed to model continuous cell-state transitions based on single-cell RNA-seq data. While mapping pseudotime to physical space is straightforward, the resulting trajectory typically does not preserve spatial relationships. Recent methods have been developed to improve spatial continuity by incorporating spatial information into the pseudotime analysis framework. In stLearn^14^, this improvement is achieved by introducing a metric that combines the pseudotime coordinate and spatial distance through a weighted average. SpaceFlow^4^ and spatialPCA^15^ integrate spatial and gene expression information via spatially-aware dimensionality reduction. However, an intrinsic difficulty in spatial trajectory analysis is balancing the conflicting demands of cell-state and spatial continuity. In complex tissues, cells that are physically proximate often exhibit distinct properties. The intrinsic incompatibility between cell-state and spatial continuity highlights the need for a new modeling framework.

We introduce **<u>O</u>**rdered **<u>N</u>**iche **<u>Tra</u>**jectory **<u>C</u>**onstruction (ONTraC for short) as a new modeling framework for analyzing spatially continuous variations in tissue microenvironments. This is achieved by disentangling environmental and cellular level changes and constructing spatial trajectories that reflect variations in the tissue microenvironment. By applying ONTraC to both simulated and real spatial transcriptomics datasets, we demonstrate its utility in characterizing continuous variation in tissue structure and in guiding downstream functional analyses.

## Results

### A Modeling Framework for Niche Trajectory Analysis

ONTraC takes spatial location and cell-type annotation information as input and uses a niche as the basic unit of the tissue microenvironment. Here, we use the term ‘niche’ to represent a multicellular, spatially localized region where different cell types may coexist and interact. In the remainder of the paper, a niche is defined as the multi-cellular region that contains the k-nearest neighbors of any cell, referred to as the anchoring cell of the niche, based on the spatial distance. The relationship between a cell and a niche is quantified by a cell-niche association score, whose value decreases with the distance from the anchoring cell. The overall property of a niche is quantified by a numerical vector summarizing its cell type composition. While neighboring cells may have distinct cellular states, the values of the cell-type composition vectors are spatially continuous, provided that k is sufficiently large.

The main output of ONTraC is the niche trajectory (**NT**), which can be viewed as a one-dimensional representation of the tissue microenvironment continuum. As schematically shown in **Fig. 1a and Extended Data Fig. 1**, the ONTraC workflow for constructing NT includes the following steps. First, we construct a niche network connecting neighboring niches, and the properties of each niche are summarized by its corresponding cell-type composition vector. Second, we construct a two-layer graph convolutional network (GCN) mode^l16^ to encode input data into low-dimensional feature vectors while maintaining spatial continuity. Third, we adopt a graph pooling network with modified loss functions^17^ to identify niche clusters and construct a niche cluster network. Fourth, we use the niche cluster network as the backbone to construct NT and determine the mapped location for each niche, which is called its NT score. Based on the cell-niche association scores, we also map each cell onto a specific location along the NT and refer to it as the cell-level NT score. The technical details are described in the **Methods** section.

**Fig. 1.**
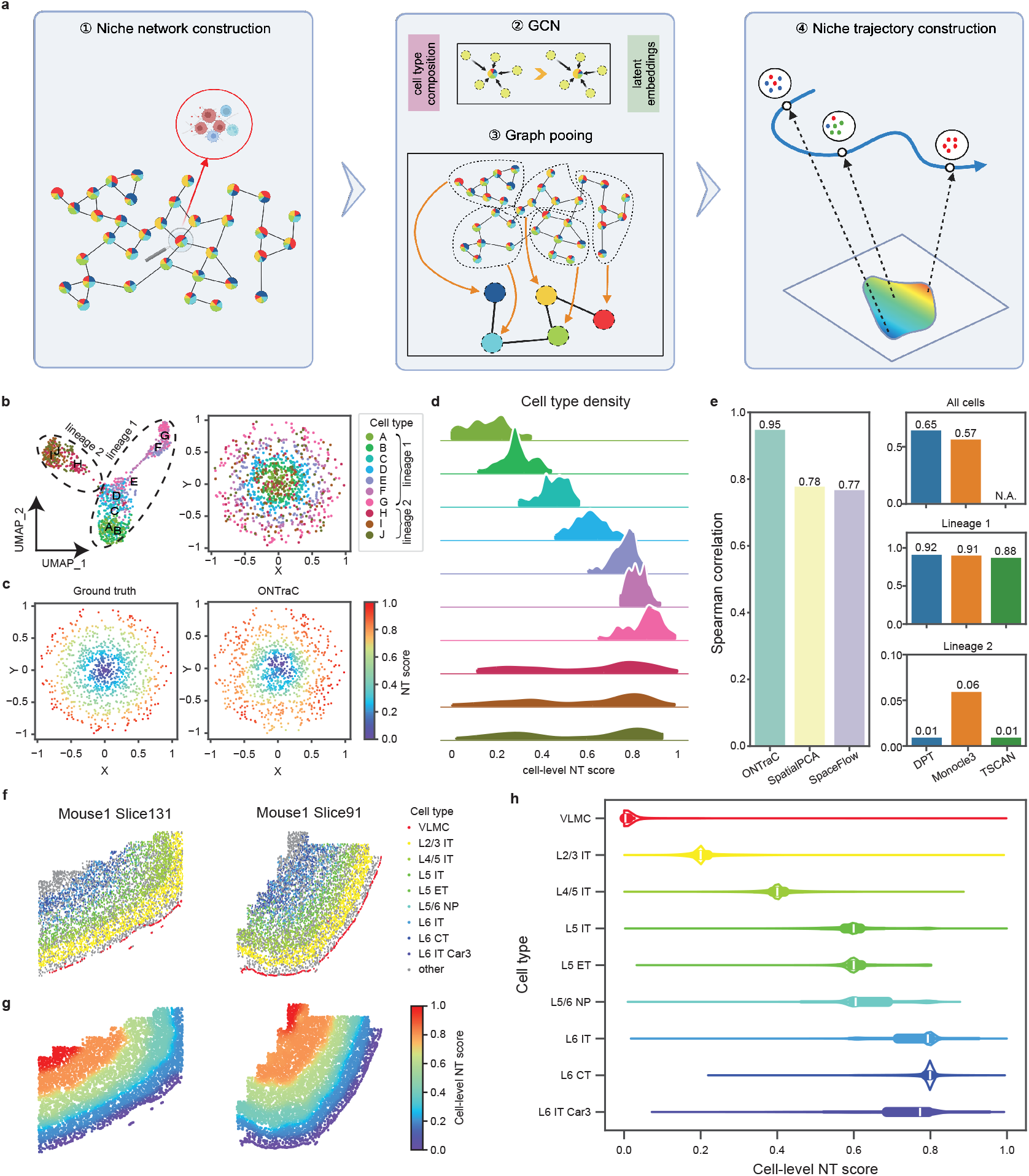
ONTraC captures spatially continuous changes in the tissue microenvironment. **a**, A schematic of the ONTraC workflow. **b-e**, Analysis of the simulated data. **b**, UMAP (left) and spatial (right) distribution of cell types in the simulated dataset. **c**, Spatial distribution of ground truth (left) and ONTraC output (right) cell-level NT scores. **d**, Cell type density along the reconstructed niche trajectory. **e**, Bar plots showing the Spearman correlation between the ground truth and the output from each method in the benchmark analysis. ONTraC, SpatialPCA and SpaceFlow were applied only to the whole dataset, whereas the other methods were applied to whole dataset, lineage 1, and lineage 2, respectively. **f-h**, Analysis of the mouse cortex MERFISH data. **f**. Spatial distribution of cell types on representative slices. **g**, Spatial distribution of cell-level NT scores on representative slices. **h**, Cell type density along the reconstructed niche trajectory.

To evaluate the performance of ONTraC, we applied it to a simulated dataset generated by superimposing spatial information over a state-of-the-art single-cell RNA-seq data simulator^18^. To mimic the generic complexity of tissue structure, we created a simulated dataset that contained multiple cell types and lineages, with only a subset of cell types exhibiting regular spatial patterns (**Methods, Supplementary Table 1 and 2**). This dataset contains the expression levels of 135 genes in 1,000 cells sampled from 10 cell types, representing two cell lineages that were separated by a bifurcation point. While cells from lineage 1 are distributed with a circular pattern in progressive order, cells from lineage 2 are distributed randomly in space (**Fig. 1b**). Applying ONTraC to this simulated dataset correctly recapitulated the spatial pattern of the niche organizations (**Fig. 1c,d, Extended Data Fig. 2**). The overall agreement between cell-level NT scores and the ground truth is high (**Fig. 1e**, Spearman correlation = 0.95).

We benchmarked the performance of ONTraC against three established pseudotime methods, including destiny^10^, Monocle 3^13^, and TSCAN^19^. For each method, we mapped the derived pseudotime coordinates to the corresponding spatial locations and evaluated their performance using Spearman correlation. For DPT and Monocle 3, the Spearman correlation values were 0.65 and 0.57, respectively, whereas for TSCAN, the overall Spearman correlation could not be evaluated because a significant fraction (over 20%) of cells were not assigned any pseudotime values (**Fig. 1e** and **Extended Data Fig. 2**). Closer examination suggested that the main challenge for these pseudotime time methods was the lack of spatial structure in lineage 2 cells. Since pseudotime analysis is often carried out on subsets of cells along a common lineage, we further tested the performance of each pseudotime analysis using cells from each lineage separately. All methods worked well for lineage 1 (Spearman correlation between 0.88 and 0.92) but performed poorly for lineage 2 (Spearman correlation between 0.01 and 0.06). Therefore, the performance of a pseudotime analysis method strongly depends on the degree of association between cell lineages and spatial locations. On the other hand, ONTraC is designed to include all the cells and its performance is robust against the presence of cell types that do not have any spatial patterns.

Furthermore, we benchmarked ONTraC against two recently developed methods, SpaceFlow^4^ and spatialPCA^15^, that extend the pseudotime framework by incorporating spatial cell-neighborhood information for cell-state embedding. To fully preserve the spatial cell-neighborhood information, we applied these methods only to the whole dataset (**Fig. 1e** and **Extended Data Fig. 2**). Both methods performed better than the pseudotime methods above, with Spearman correlation at 0.77 and 0.78, respectively (**Fig. 1e**), suggesting that incorporating spatial information is beneficial for spatial trajectory analysis. However, they still cannot correctly map lineage 2 cells onto the spatial trajectory (**Extended Data Fig. 2)**. Taken together, these analyses indicate that ONTraC is more accurate than existing methods in spatial trajectory reconstruction.

### Dissecting the spatial structure of the mouse motor cortex using niche trajectory analysis

We applied ONTraC to analyze a MERFISH dataset from the mouse motor cortex, which includes 61 tissue slices and contains approximately 280,000 cells characterized by a 258-gene panel^20^. The study identified 23 transcriptional subclasses, several of which exhibit distinct spatial patterns (**Fig. 1f**). Interneurons are distributed across all cortical layers, with each associated with a distinct IT subclass. The vascular leptomeningeal cells (VLMCs) demarcate a thin layer at the border of the cortex. We ran the ONTraC analysis treating each subclass as a ‘cell type’, while aggregating the remaining cells under the cell type ‘other’. Using information from all 24 cell types (i.e., 23 subclasses + other), we constructed the niche trajectory by applying the ONTraC workflow. The resulting NT faithfully reveals the underlying layer structure (**Fig. 1g**), starting from the outer layer and progressively moving inward. The sharp transition between the layer boundaries is also preserved. The cell-type composition varies continuously along the NT, with each layer-specific cell type enriched within a narrow section (**Fig. 1h**). On the other hand, non-neuronal cell types such as astrocytes and microglia spread broadly (**Extended Data Fig. 3a**).

In the original study^20^, the authors applied pseudotime analysis to order IT neurons. We compared the cell-level NT scores obtained from ONTraC with the pseudotime values for IT neurons and found that the results from these two approaches are highly correlated (R = 0.92) (**Extended Data Fig. 3b**). However, similar to our simulation analysis above, pseudotime analysis does not work well if this analysis was based on all the cells ro another subset. Therefore, the results from pseudotime analysis are subject to prior knowledge biases, whereas ONTraC does not suffer from this limitation.

As an additional validation, we tested whether the cell-level NT scores correlate with cortical depths. A similar analysis was performed in the original paper and resulted in high correlation ^20^, but the analysis was limited to IT neurons. In contrast, our ONTraC included all the cells (see **Methods** for details). We observed a high degree of correlation between the cell-level NT scores and cortical depths (**Extended Data Fig. 3c**). For over 80% (49/61) of the tissue sections, the Spearman correlation between cell-level NT scores and cortical depths is 0.90 or higher. However, a small number of samples exhibited significantly lower correlations. For example, in the Mouse1 Slice131 sample, the correlation was only 0.80. To understand the underlying cause, we compared the cell-type distribution between the Mouse1 Slice131 and Mouse1 Slice91 samples. In the latter sample, the Spearman correlation was 0.99. We noticed that the VLMC layer is completely included in Mouse1 Slice91 but is partially missing in Mouse1 Slice131 (**Fig. 1f**). The discrepancies between cell-level NT scores and cortical depths are caused by the error in the cortical depth calculation method, which is sensitive to the missing VLMC layer information (**Extended Data Fig. 3c, insets)**. On the other hand, our ONTraC method is reference free, thereby providing a more robust approach for evaluating cortical depth.

### ONTraC analysis identifies tissue microenvironment associated cell-state variations in developing dorsal midbrain

Tissue microenvironment plays a distinctly important role during development, where cells that are initially identical undergo divergent paths during differentiation. To investigate the role of the tissue microenvironment in mediating cell state changes, we analyzed a public stereo-seq dataset from mouse embryo development^21^. This dataset includes whole-embryo, transcriptome-wide, single-cell gene expression profiles across eight time points between E9.5 and E16.5. We focused on a subset that was analyzed extensively in the original study, containing over 26,000 segmented cells from the dorsal midbrain region across three time points: E12.5, E14.5, and E16.5. We annotated cell types following the same procedure as in the original study^21^ (see **Methods**). Consistent with the original study, we identified ten cell types, including radial glial cell (RGC), glioblast (GioB), neuroblast (NeuB), glutamatergic neuroblast (GluNeuB), glutamatergic neuron (GluNeu), GABAergic neuron (GABA), Basal, Fibroblast (Fibro), Endothelial (Endo), and Erythroid (Ery) cells (**Extended Data Fig. 4**). While the original study also detected a small cell cluster annotated as microglia, our analysis did not identify such a cluster. This discrepancy is likely due to the intrinsic uncertainty associated with rare cell type detection.

To identify changes in tissue structure across developmental stages, we trained a common ONTraC model for all the samples, ensuring that the resulting NTs are directly comparable. Within each sample, the NT progresses continuously from caudal to rostral and from ventral to dorsal areas (**Fig. 2a, Extended Data Fig. 5**). Along the niche trajectory, the most enriched cell type transitions from RGCs to increasingly differentiated cell types (**Fig. 2b, Extended Data Fig. 6**). Comparing the NT score distributions across different samples, we noted a significant increase as development progresses (**Extended Data Fig. 7**). Altogether, these results suggest that the niche trajectory delineates a path from an undifferentiated to increasingly more differentiated tissue microenvironment.

**Fig. 2.**
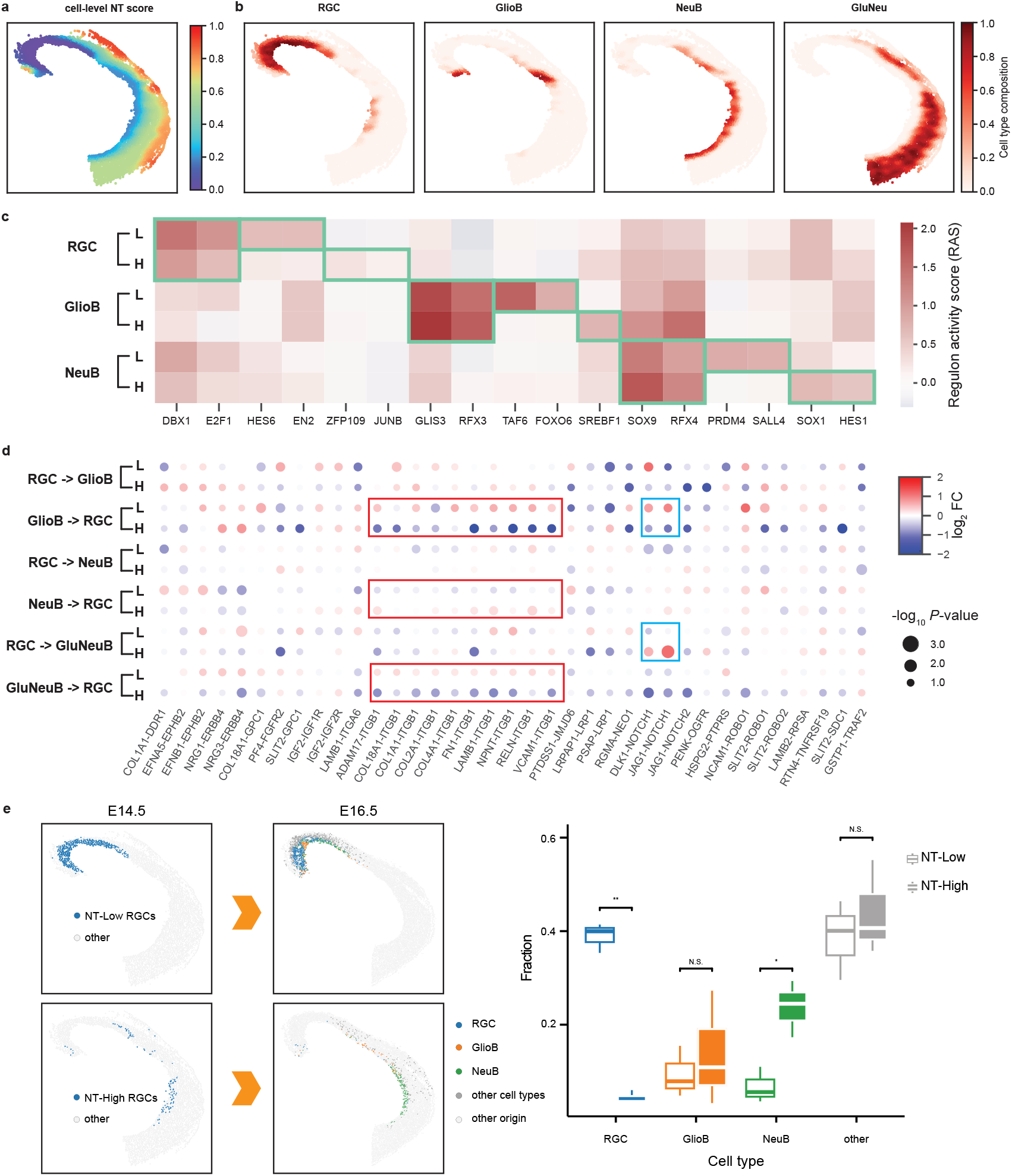
ONTraC analysis reveals tissue-microenvironment-mediated cell-state changes in E14.5 mouse dorsal midbrain. **a**, Spatial distribution of cell-level NT scores. **b**, Spatial distribution of representative cell types. c, Changes in regulon activity associated with cell type and microenvironment. Green boxes highlight cell-type or microenvironment-specific high regulon activities. (H: NT-High; L: NT-Low). **d**, Cell-type-pair dependent ligand-receptor activities in NT-High (H) and NT-Low (L) microenvironments. ITGB1 and NOTCH1 associated signals are highlighted by red and blue boxes, respectively. **e**, Moscot analysis mapping E14.5 NT-Low and NT-High RGCs to putative offspring cells in E16.5. Left: spatial locations of original and mapped cells; right: Box plots showing the cell-type composition of putative offspring cells of E14.5 NT-Low and NT-High RGCs. Statistics are based on all three independent E16.5 mouse samples. N.S. (non-significant); * (p-value < 0.05); ** (p-value < 0.01).

Among these cell types, RGCs are especially notable due to their ability to migrate and differentiate into either neuronal or glial cells during embryonic development. The spatial distribution of RGCs exhibits distinct spatiotemporal patterns (**Fig. 2b** and **Extended Data Fig. 6**). At E12.5, RGCs occupy nearly the entire dorsal midbrain but become increasingly spatially restricted during development. At E14.5, they are distributed along a narrow streak at the ventral edge. By E16.5, the RGCs are confined to the caudal domains.

To investigate whether changes in the tissue microenvironment play a role in mediating the gene expression programs in RGCs, we focused on the E14.5 time point, when the RGCs are distributed across a wide range along the niche trajectory (**Fig. 2a, b**), indicating exposure to diverse tissue microenvironments. We aimed to determine if such diversity could lead to differential transcriptional responses and, if so, whether these changes are associated with cell-fate decisions during development. To this end, we divided the E14.5 RGCs into two subgroups based on their cell-level NT scores: NT-Low (< 0.17) and NT-High (≥ 0.17). By comparing the gene expression levels of these two RGC subgroups, we identified 34 differentially expressed genes (DEGs), including 12 up-regulated and 22 down-regulated genes (**Extended Data Fig. 8a** and **Supplementary Table 3**). Among these DEGs, several have been previously shown to play functional roles in neurogenesis, such as Efna5^22^, Ccnd2<u>^23^</u>, and Tet3^24^. Through gene set enrichment analysis (GSEA)^25^, we found several significantly enriched Gene Ontology (GO) terms associated with neurogenesis, such as NEUROEPITHELIAL_CELL_DIFFERENTIATION (NES = 1.80, *P*-value = 0.0026), and NEGATIVE_REGULATION_OF_NERVOUS_SYSTEM_DEVELOPMENT (NES = 1.78, *P*-value = 0.0020) (**Extended Data Fig. 8b**). The level of statistical significance is moderate, potentially due to the small number of genes analyzed.

To gain mechanistic insights, we performed gene regulatory network (GRN) analysis using the SCENIC workflow^26^. Consistent with the original study^21^, our analysis identified several known cell-type-specific regulators, such as DBX1 and E2F1 for RGCs, GLIS3 and RFX3 for glioblasts, and SOX9 and RFX4 for neuroblasts (**Fig. 2c and Extended Data Fig. 9**). Whereas the original study only examined GRN differences between cell types, here we were able to further investigate within cell-type variations by leveraging the NT structure. By comparing the RAS scores associated with NT-Low and NT-High subgroups, we identified several regulons whose activities change significantly along the NT, such as HES6 and EN2 (in RGCs), FOXO6 and SERBF1 (in GlioB), and PRDM4 and SOX1 (in NeuB) (**Fig. 2c**). This suggests that these regulators may play a role in fine-tuning the timing of cell differentiation in a niche-dependent manner.

Next, we performed cell-cell interaction analysis using the Giotto pipeline<u>^27,28^</u>, and compared the patterns between NT-Low and NT-High niches (see **Methods**). Cell proximity analysis indicates that RGCs tend to be isolated from other cell types in NT-Low niches, whereas in NT-High niches, they are more likely to interact with glioblasts, neuroblasts, and glutamatergic neuroblasts (**Extended Data Fig. 10**). Furthermore, ligand-receptor pair analysis identified significant changes between NT-Low and NT-High niches (**Fig. 2d**) associated with NOTCH1 and Integrin beta 1 (ITGB1) signaling pathways, both are known to regulate radial glia differentiation<u>^29,30^</u>.

To further investigate the connection between NT and cell fate changes, we linked cells obtained from different time points using the Moscot procedure^31^ (see **Methods**). We compared the predicted offspring associated with NT-Low and NT-High RGCs and observed significant differences (**Fig. 2e**). Whereas the NT-Low RGC offspring cells tend to retain the RGC identity, the NT-High RGC offspring cells are more differentiated (**Fig. 2e**). Taken together, the above analyses strongly suggest that ONTraC is a useful tool for dissecting the role of tissue microenvironment changes in mediating transcriptional changes and cell-fate decisions.

## Discussion

We have developed ONTraC as a new framework for constructing spatial trajectories. This framework is generally applicable, as it requires only cell-type annotation and spatial information as inputs. Such information can be obtained not only from spatial transcriptomics data but also from other data modalities. A key feature of ONTraC is that it treats a niche, rather than a cell, as the basic unit. While the use of niche structure in spatial analysis is not new^7,32^, ONTraC differs from previous approaches in that it explicitly models the hierarchy of niche-level and cell-level properties. Specifically, the niche-level properties are shared among all cells within a niche, rather than being limited to any specific cell type or cell state. On the other hand, cell-level properties are maintained and can be compared at the single-cell resolution. This strategy naturally incorporates local cellular heterogeneity as a source of spatial variation at the niche-level, while simultaneously providing a common framework to study coordinated responses from multiple cell types within a tissue microenvironment. Through analysis of both simulated and real spatial transcriptomics datasets, we have shown that ONTraC is not only effective for detecting continuous tissue microenvironment variations, but also provides a useful modeling framework to study impact of the tissue-environment changes on mediating cell-state dynamics.

Inspired by previous work from several groups^2,4,7^, ONTraC utilizes a GNN modeling framework to integrate spatial and cell-type composition information. In previous work, the use of GNN was limited to characterizing spatial domains, but here we have extended its use for constructing spatial trajectories. Spatial information is preserved throughout the ONTraC workflow, as all the analyses are supported by the underlying spatial network structure. The loss function is designed to properly balance the conflicting demands for preserving both node property similarity and spatial continuity. Our analyses show that ONTraC performs well in analyzing real biological datasets in various contexts.

ONTraC has several limitations. First, since it requires cell-type annotation information as input, ONTraC is designed primarily for analyzing spatial data with single-cell resolution. On the other hand, it is possible to adapt the ONTraC workflow for the analysis of lower-resolution data, for example, by using either spot-level clustering or cell-type deconvolution results as a proxy for cell-type annotation. Second, the outcome of ONTraC for a given spatial omic dataset depends upon how the cell types are annotated, therefore different choices of clustering methods and/or parameter settings could lead to variations. Third, the output of ONTraC is an unbranched trajectory, which may be over-simplified for representing the spatial structure of a complex tissue. Future extensions are needed to overcome these limitations.

Finally, there is a growing interest in integrating spatial omic and pathological information to study human diseases^33^. Such integrations could lead to comprehensive understandings of the disease progression process and further provide mechanistic insights. Niche trajectory analysis may provide a useful tool for such integrations.

## Methods

### The ONTraC workflow

The ONTraC workflow includes the following steps: 1. Constructing a niche network; 2. Using GCN to encode input data in a low-dimensional feature space; 3. Using graph pooling to identify niche clusters and their spatial relationship; 4. Finalizing niche trajectory construction and NT-score evaluation (**Fig. 1a** and **Extended Data Fig. 1**). The details are described in the following.

#### Step 1: Constructing a niche network

A niche is defined as a multicellular, spatially localized region where different cell types may coexist and interact. Although the general workflow is sufficiently flexible to accommodate variations in the precise definition of niches, the analyses in this paper were conducted using the following procedure to ensure reproducibility: Each niche is anchored at a single cell, which is referred to as the anchoring cell, and including its *k*-nearest neighbors (kNN) (default *k* = 50) according to physical distance. A niche network is created by connecting pair of niches whose anchoring cells are mutual nearest neighbors. The edge relationship in the niche network is represented by the binary adjacency matrix *A* = (*a*_*ij*_).

Since neighboring niches may overlap, each cell is typically included in multiple niches. The degree of association between a cell *i* and a niche *j* that contains it is quantified by the cell-niche association score, which is defined by 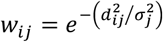, where *d*_*ij*_ denotes the physical distance between the cell *i* and the anchoring cell of niche *j*, and σ_*j*_ represents a niche-specific normalizing factor. By default, the value of σ_*j*_ is set to be the spatial distance between the anchoring cell and its *20*th nearest neighbor. Based on its cell-type composition, a niche *j* is assigned with a numerical vector **v**_***j***_ = (*v*_*jm*_), whose values are given by *v*_*jm*_ = ∑_*i*_ *w*_*ij*_*I*_*im*_/∑_*i*_ *w*_*ij*_, where *I*_*im*_ is a binary indicator of whether the cell *i* belongs to the cell-type *m*.

#### Step 2: Using GCN to encode input data in a low-dimensional feature space

We use a two-layer GCN model to integrate spatial and cell-type composition information, which can be mathematically represented as:

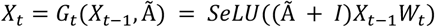

where *X*_*t*_ (t = 1 or 2), represents the low-dimensional embedding vectors (dimension = 4 by default) at the *t*-th layer, while *X*_o_ represents the input cell-type composition vectors **v**_***j***_ obtained from Step 1.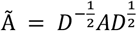 represents the normalized adjacency matrix, with *D* = *diag*(*A*), and *W*_*t*_ represents the trainable parameters. The model training process is described later.

#### Step 3: Using graph pooling to identify niche clusters and their spatial relationship

In this step, we use a modified graph pooling approach^17^ to convert the original spatial network into a network of niche clusters. To this end, we probabilistically cluster niches based on the embedding vector values generated in Step 2. The main output is a matrix *C* = (*c*_*jk*_), whose values *c*_*jk*_ represent the probabilistic assignment of a niche *j* into cluster *k*. For this purpose, we use a linear neural network model, *C* = softmax (*βX*_2_*W*_*c*_), where *β* is a tuning parameter set to a default value of 0.03, and *W*_*c*_ contains trainable parameters.

Next, we perform graph pooling by collapsing the niche network to a niche cluster network. For each pair of niche clusters *k* and *l*, the corresponding edge connectivity is given by *E*(*k, l*) = ∑_*i*_ ∑_*j*_ *c*_*ik*_*a*_*ij*_*c*_*jl*_, where (*a*_*ij*_) and (*c*_*jk*_) represent the adjacency matrix and the niche cluster assignment matrix, respectively. The edge connectivity is further normalized by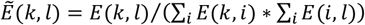.

#### Step 4: Finalizing niche trajectory construction and NT-score evaluation

In the following, we use *N, M*, and *K* to represent the total number of niches, cell types, and niche clusters, respectively. To construct the niche trajectory, we search for the optimal ordering of niche clusters: *p*_1_ → *p*_2_ → … → *p*_*k*_ that maximizes the total edge connectivity, which is defined by 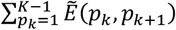. Then, the niche cluster *p*_*k*_ is assigned a score *s*(*p*_*k*_) = (*k* − 1)/(*K* − 1). Next, we map each niche *j* along the trajectory by considering its probabilistic assignment to each niche cluster. The corresponding coordinate, called the **NT score**, is defined as:

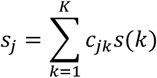

The range of NT scores is between 0 and 1. Finally, to account for the varying degrees of association between cells and niches, we further define the **cell-level NT score** for each cell *i* as follows:

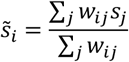

The cell-level NT score is used to stratify cells to study within cell-type variations mediated by tissue microenvironment changes.

#### Model training

The ONTraC workflow contains two sets of trainable parameters *W*_*t*_ (Step 2) and *W*_*c*_ (Step 3). These parameters are trained to optimize spatial and cell-type composition continuity. Towards this end, we design a loss function that contains three terms representing modularity, purity, and regularization.

The modularity loss term is given by:

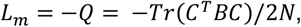

where *Q* is known as the modularity ^34^, and *C* is the probabilistic cluster assignment matrix defined in Step 3. *B* is the modularity matrix, which is defined by 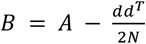, where *d* is the node degree vector. This term is included to enhance modularity.

The purity loss term is given by:

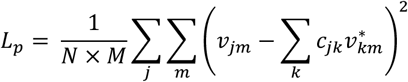

where **v**_***j***_ = (*v*_*jm*_) represents the cell-type composition vector associated with niche *j*, and 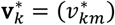 represents the average cell-type composition vector within niche cluster *k*. This term is introduced to enhance feature similarity within each niche cluster.

Finally, the regularization loss term is given by:

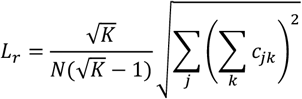

This term is introduced to regularize the distribution of cluster sizes ^17^.

Taken together, the overall loss function is given by

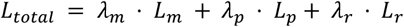

For the analyses in this paper, we use the following parameter setting: *λ*_*m*_ = 0.3, *λ*_*p*_ = 300, and *λ*_*r*_ = 0.1. The values of the trainable parameters *W*_*t*_ and *W*_*c*_ are obtained using Adam optimizer^35^ with 0.03 as the learning rate. The model is implemented using PyTorch (https://pytorch.org/) and PyTorch Geometric (https://github.com/pyg-team/pytorch_geometric), and carried out on a NVIDIA A100 GPU. In our analysis, we set the batch size to 5 (stereo-seq dataset) and 10 (MERFISH dataset) with a maximum of 1000 epochs.

### Simulated time generation

Simulated single-cell gene expression data were generated using *dyngen*^18^ with the default bifurcating backbone, which contains two cell lineages diverging from a single bifurcation point. A total of 1000 cells were sampled in 10 batches. Based on their relative positions along the trajectory, the cells were assigned to 10 different cell-types (A-J), with cell types A-G mapped to lineage 1 and cell types H-J to lineage 2. The simulated gene expression profiles contain 135 genes, including 35 transcription factors, 50 target genes, and 50 housekeeping genes.

Next, each cell is assigned to a spatial location as follows: Each cell *i* in lineage 1 is mapped to a location (*x*_*i*_, *y*_*i*_), which is randomly chosen from the circle 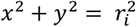. The radius *r*_*i*_ is proportional to the simulated time *t*_*i*_ obtained from *dyngen* output, and rescaled so that its maximum value is 1. Each cell in lineage 2 is mapped to a random location in the area defined by *x*^2^ + *y*^2^ ≤ 1. As the niche-level, the spatial trajectory progresses radially from the center of the circle, and the ground-truth cell-level NT score is equal to *r*_*i*_ for all cells. While the ordering of lineage 1 cells along the spatial trajectory maintains their lineage relationship, there spatial distribution of lineage 2 cells is unrelated to their lineage relationship.

### Benchmark analysis

To benchmark the performance of ONTraC, we considered five existing methods for comparison and applied each method to the simulated dataset, including three pseudotime analysis methods: destiny (v3.16.0)^10^, Monocle 3 (v1.3.7)^13^, and TSCAN (v2.0.0)^11^, as well as two methods that further incorporate spatial information: SpaceFlow (v1.0.4)^4^ and SpatialPCA (v1.3.0)^15^. For each pseudotime analysis method, three sets of results were generated, corresponding to all cells, lineage 1 cells, and lineage 2 cells, respectively. On the other hand, SpaceFlow and SpatialPCA were applied only to analyze all cells to faithfully represent the spatial information.

For destiny, dimensionality reduction was performed using ‘*DiffusionMap*’ with default parameters, and pseudotime values were generated using the ‘*DPT*’ function. For Monocle 3, dimensionality reduction was achieved using the ‘*reduce_dimension*’ function with the UMAP method, and pseudotime values were obtained by using the ‘*pseudotime*’ function. The root cell was selected to be the true starting cell along the trajectory. For TSCAN, pseudotime values were obtained by sequentially applying the ‘*preprocess*’, ‘*exprmclust*’, and ‘*TSCANorder*’ functions.

For SpaceFlow, pseudotime values were obtained by sequentially applying the ‘*preprocessing_data*’, ‘*train*’, ‘*segmentation*’, and ‘*pseudo_Spatiotemporal_Map*’ methods with default parameters. For Spatial PCA, spatial principal component analysis was performed by sequentially applying the ‘*SpatialPCA_buildKernel*’, ‘*SpatialPCA_EstimateLoading*’, and ‘*SpatialPCA_SpatialPCs*’ functions. Cell clusters were obtained using the ‘*walktrap_clustering*’ function with parameters “clusternum=6, knearest=32”. The pseudotime was obtained using Slingshot (v2.10.0) following the tutorial of SpatialPCA.

### Analysis of mouse cortex MERFISH data

Processed mouse motor cortex MERFISH data, including count matrix, cell type annotation, spatial location, and other meta information, were obtained from Brain Image Library (https://doi.brainimagelibrary.org/doi/10.35077/g.21). For cell type annotation, we treated each of the 23 transcriptome subclasses identified by the original study as a distinct cell type. Additionally, another cell type, termed ‘other’, was used to annotate those cells that do not fall into any of these transcriptional classes. The cortical depth is calculated as the median distance to all VLMC cells. We performed pseudotime analysis by using the destiny (v3.16.0) using the same setting as in the benchmark analysis. Only IT neurons were included in this analysis as in the original study.

### Analysis of mouse embryo dorsal midbrain stereo-seq data

Mouse embryo stereo-seq data were downloaded from the MOSTA website (https://db.cngb.org/stomics/mosta/download/). We analyzed the dorsal midbrain subset corresponding to the following five samples: E12.5 E1S3, E14.5 E1S3, E16.5 E1S3, E16.5 E2S6, and E16.5 E2S7. Segmented cell locations and the single-cell resolution gene expression profiles were generated by the authors. Because the authors did not provide cell-type annotation information, we regenerated the cell-type labels by following the same procedure as described by the authors. Briefly, we normalized gene expression using SCTransform^36^ (as implemented in Seurat v4.3.0) and removed the batch effect using harmony (v1.2.0)^37^. This was followed by dimension reduction using UMAP^38^ and Leiden clustering^39^. The identified clusters were annotated using known cell-type specific marker genes. Comparing the UMAP (**Extended Data Fig. 2**) with the original study (**Figure 5A** in ref^.21^) shows similar structures, indicating the overall cell-type structure is reproduced. The remaining discrepancy is likely due to the difference between our parameter setting and software version and the original study.

To aid the analysis of cell state variation along the NT. For the E14.5 sample, we divided cells into two subgroups by thresholding the cell-level NT scores (cutoff = 0.17). These subgroups are named NT-Low or NT-High, respectively. We used ‘FindMarkers’ in Seurat (v4.3.0) to identify DEGs between NT-Low and NT-High cells based on the two criteria: abs(FC) > 0.2 and p-value < 0.01. Gene set enrichment analysis was carried out by using the *fgsea* package (v1.28.0) in BioConductor^40^.

### Gene regulatory network analysis

Transcription factor regulons were identified using the SCENIC workflow^26^ as implemented in pySCENIC v0.12.1. A gene co-expression network was inferred using the ‘grn’ module with default parameters. Motif enrichment analysis for each gene co-expression module was predicted using the *‘ctx’* module combined with the pre-computed SCENIC+ database (mm10 refseq_r80 v10). Regulon activity scores (RAS) were quantified at the single-cell level using AUCell and then normalized by transforming them into z-scores within each developmental stage. Regulon specificity scores (RSS) were calculated using the ‘*regulon_specificity_scores*’ function based on the algorithm described in ref.^41^

### Cell-cell Interaction analysis

The Giotto Suite package^27,28^ is used for cell proximity and cell-cell interaction analyses. Spatial networks were constructed using the ‘*createSpatialNetwork*’ function with the parameter setting “*method = ‘knn’, k = 5*”. Cell proximity analysis was performed using the ‘*cellProximityEnrichment*’ function with parameter ‘*number_of_simulations = 1000*’. Ligand-receptor enrichment analysis was performed by using the ‘*exprCellCellcom*’ and ‘*spatCellCellcom*’ functions with the parameter setting “*random_iter = 1000, adjust_method = ‘fdr’* ‘‘.

### Spatiotemporal mapping of cells across different samples

The Moscot pipeline^31^, with its default parameter setting, was used for spatiotemporal mapping between E14.5 and E16.5 dorsal midbrain stereo-seq samples. The main output of Moscot is a learned coupling matrix that probabilistically connects cells from an early to a late time point. For each RGC from the E14.5 sample, we identified its putative offspring cells in the E16.5 sample as the set of five cells with the highest non-zero coupling probabilities. Depending on their cell origin, a putative offspring cell is labeled either as ‘NT-Low offspring’ or ‘NT-High offspring’. To eliminate uncertainty, the cells identified as the putative offspring of both NT-Low and NT-High RGCs are excluded from further study.

## Data and code availability

The ONTraC method is implemented as a Python package, which is available on GitHub (https://github.com/gyuanlab/ONTraC) with detailed instructions for installation, execution, and post-analysis. The simulated and example data are available in ‘examples’ directory.

## Acknowledgements

We thank Dr. Joselyn Cristina Cháavez-Fuentes for testing the ONTraC and Dr. Panos Roussos for feedback regarding an earlier version of the manuscript. This research was supported by NIH grants RF1MH133703, RF1MH128970 and R01AG066028 to G.Y. This work was supported in part through the computational and data resources and staff expertise provided by Scientific Computing and Data at the Icahn School of Medicine at Mount Sinai and supported by the Clinical and Translational Science Awards (CTSA) grant UL1TR004419 from the National Center for Advancing Translational Sciences.

## Notes

### Competing Interest Statement

The authors have declared no competing interest.

