## Supplementary figures and images for "Characterizing Spatially Continuous Variations in Tissue Microenvironment through Niche Trajectory Analysis"

### Extended Data Fig. 1

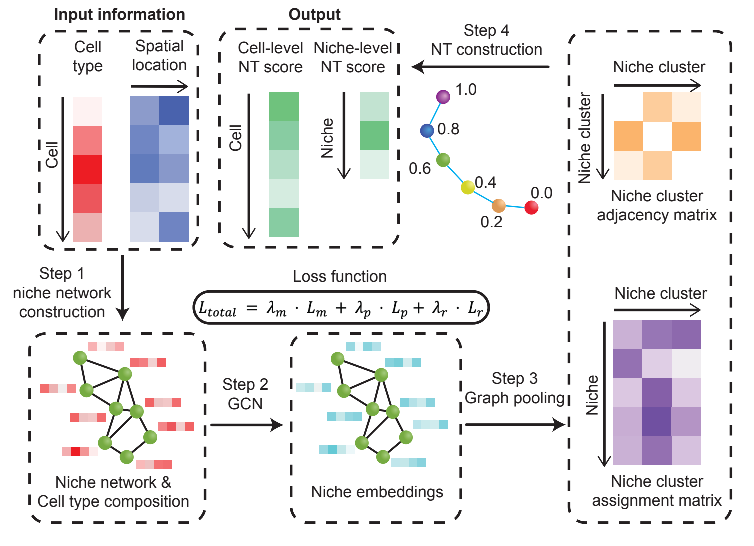

### Extended Data Fig. 2

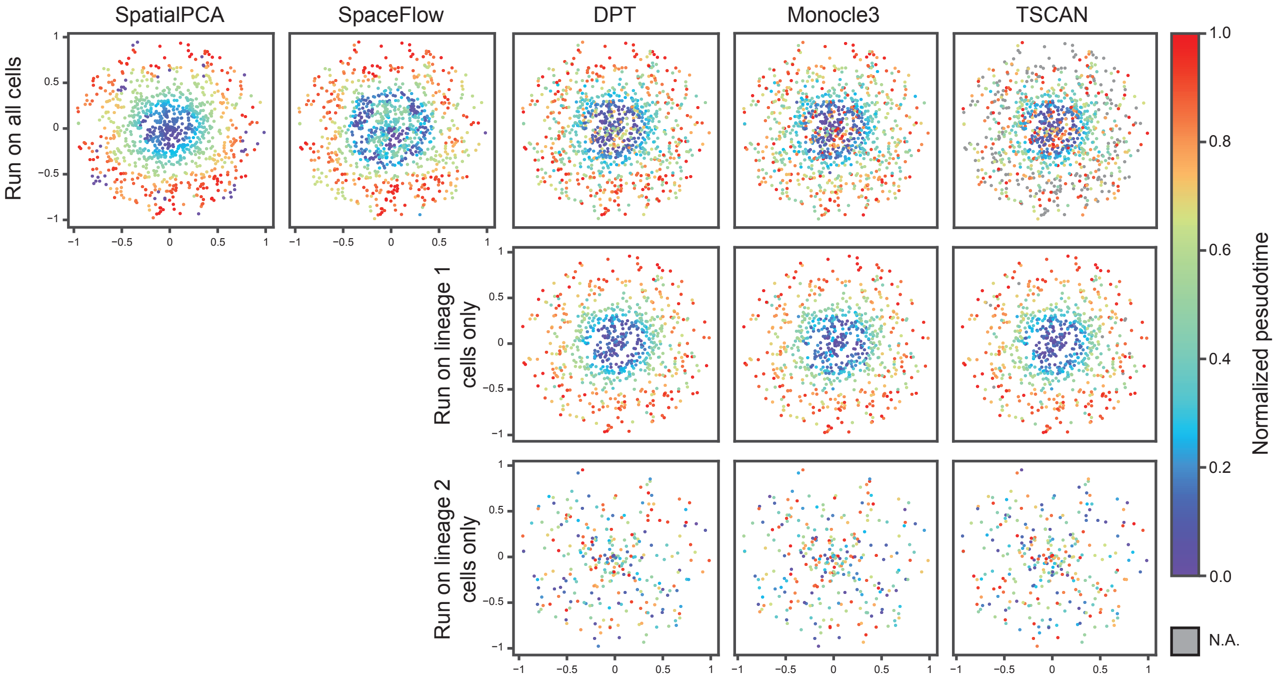

### Extended Data Fig. 3

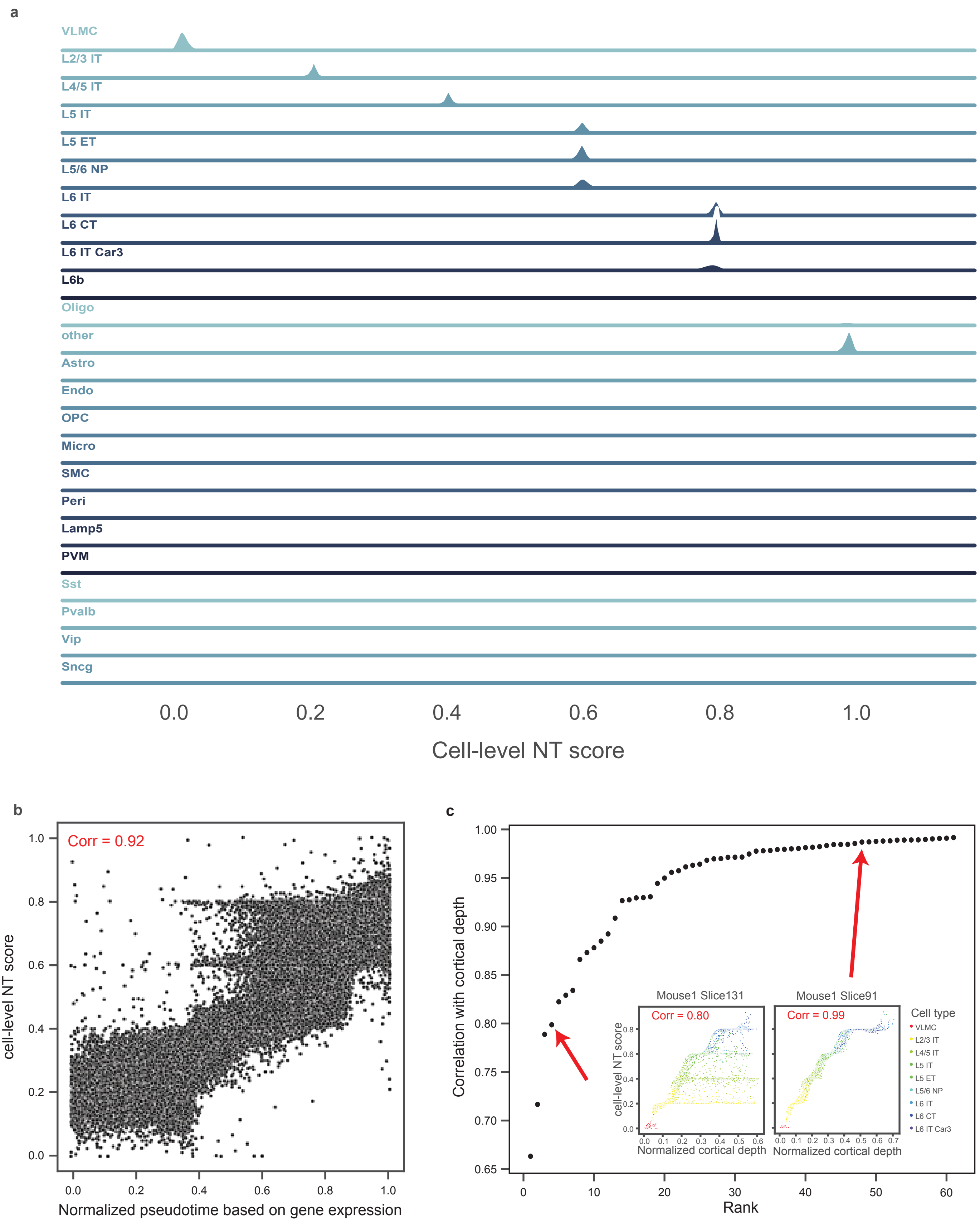

### Extended Data Fig. 4

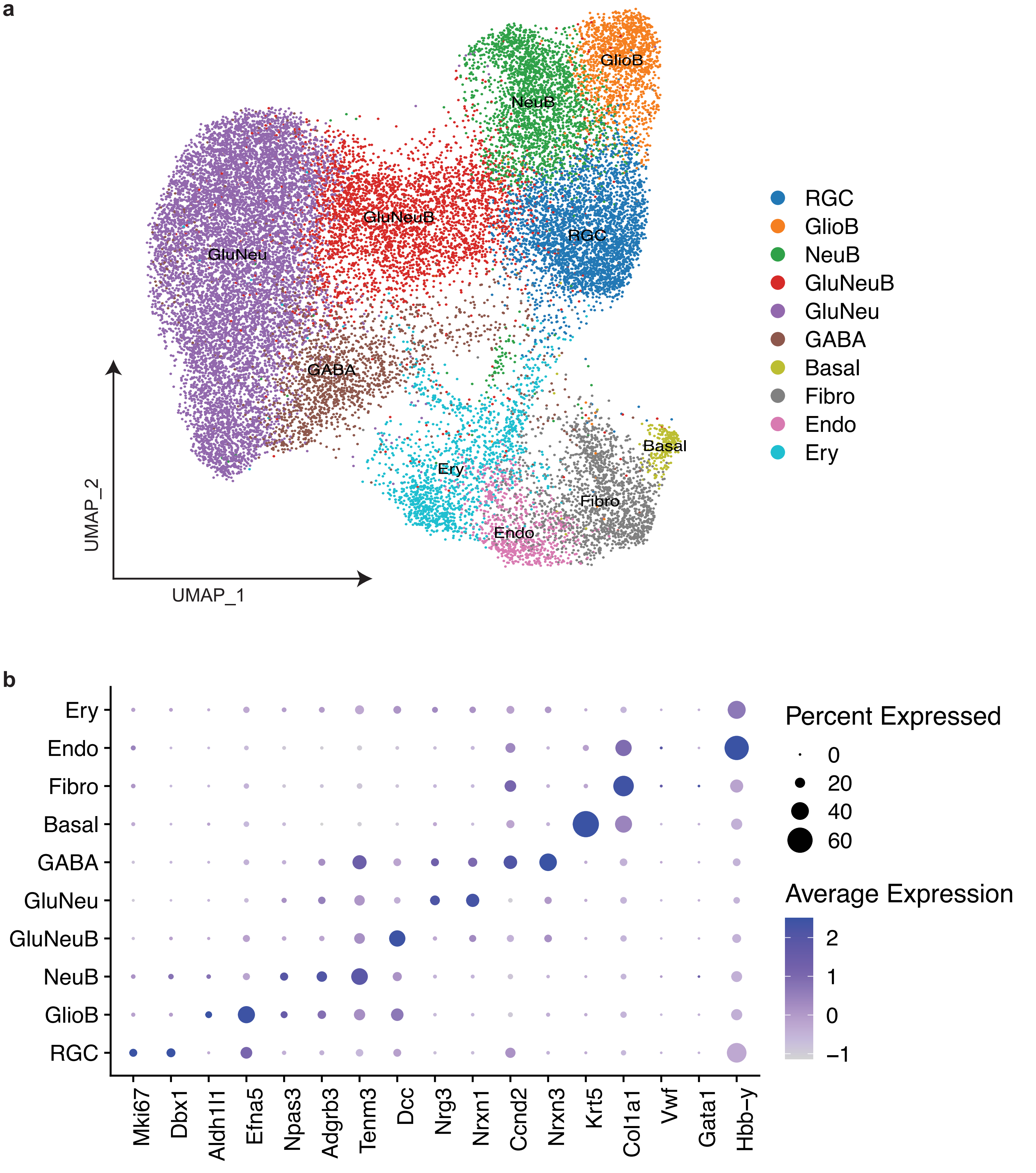

### Extended Data Fig. 5

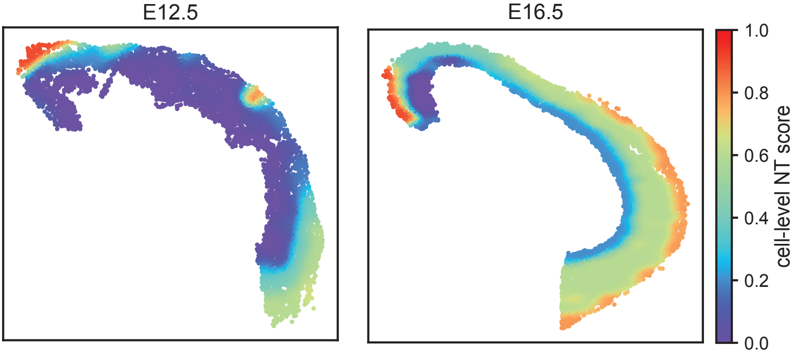

### Extended Data Fig. 6

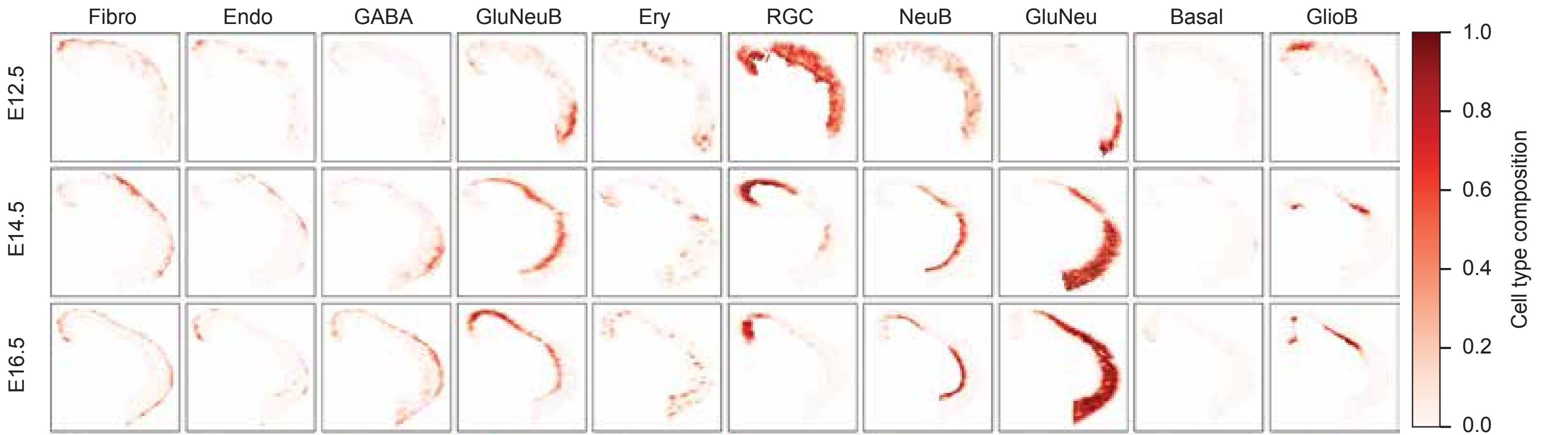

### Extended Data Fig. 7

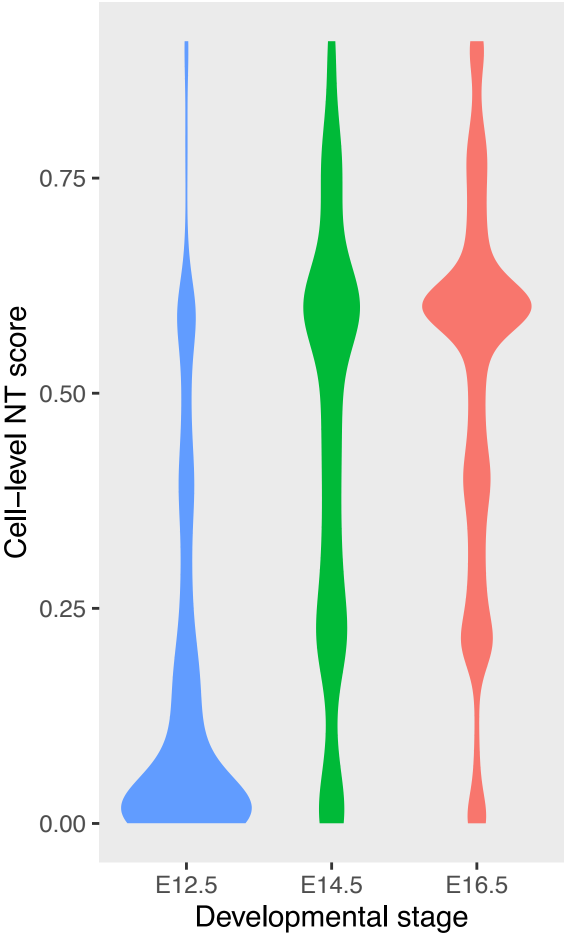

### Extended Data Fig. 8

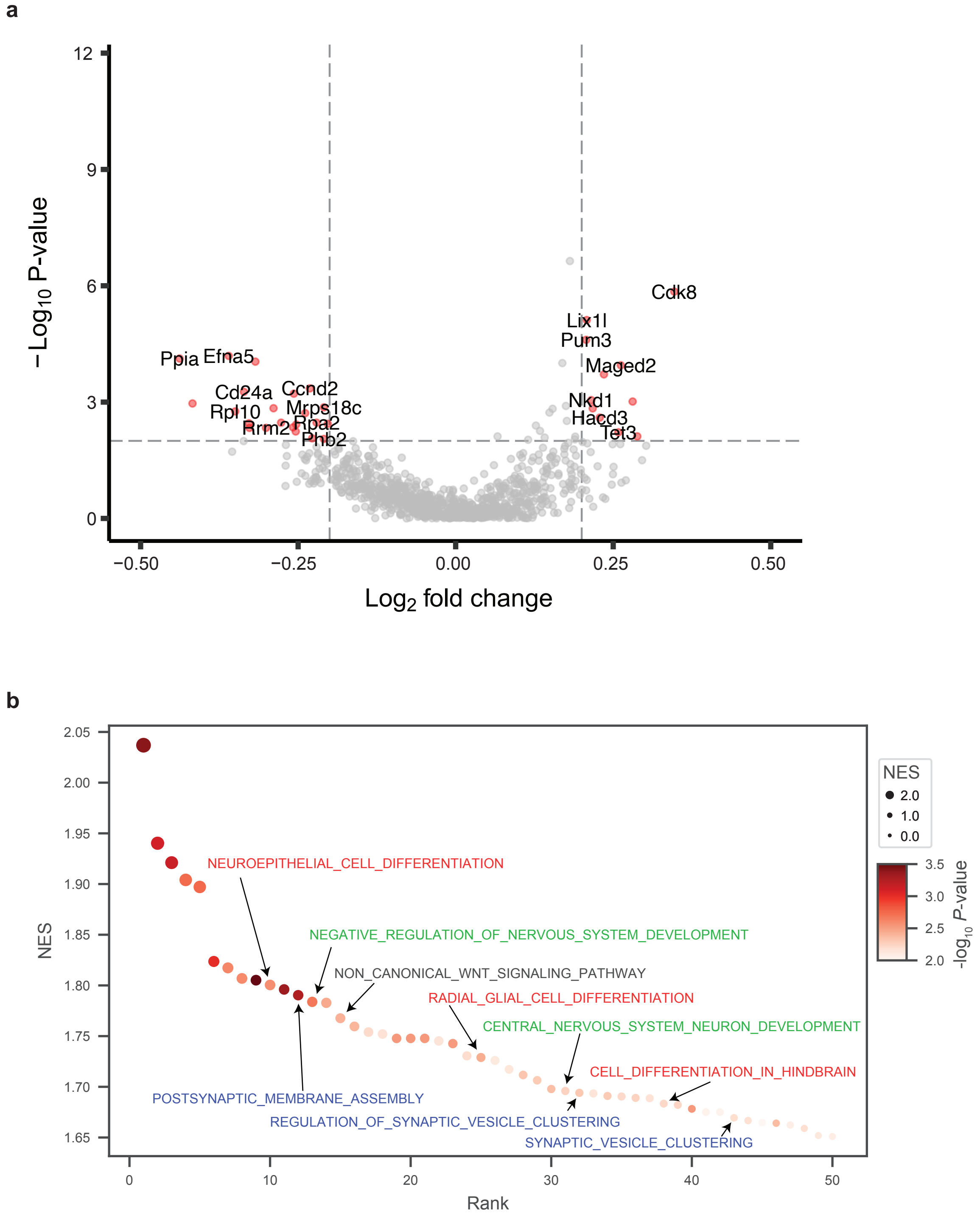

### Extended Data Fig. 9

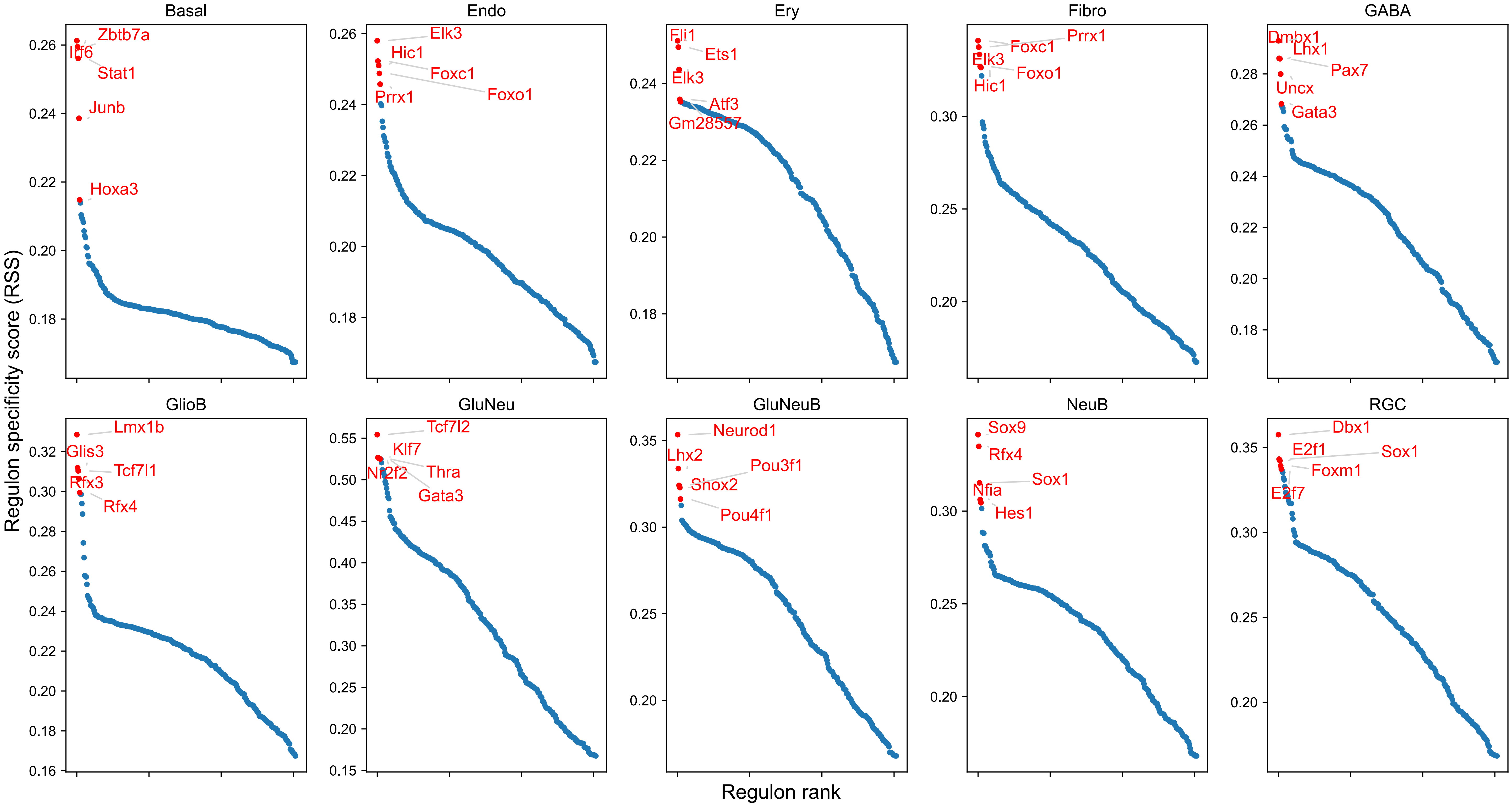

### Extended Data Fig. 10

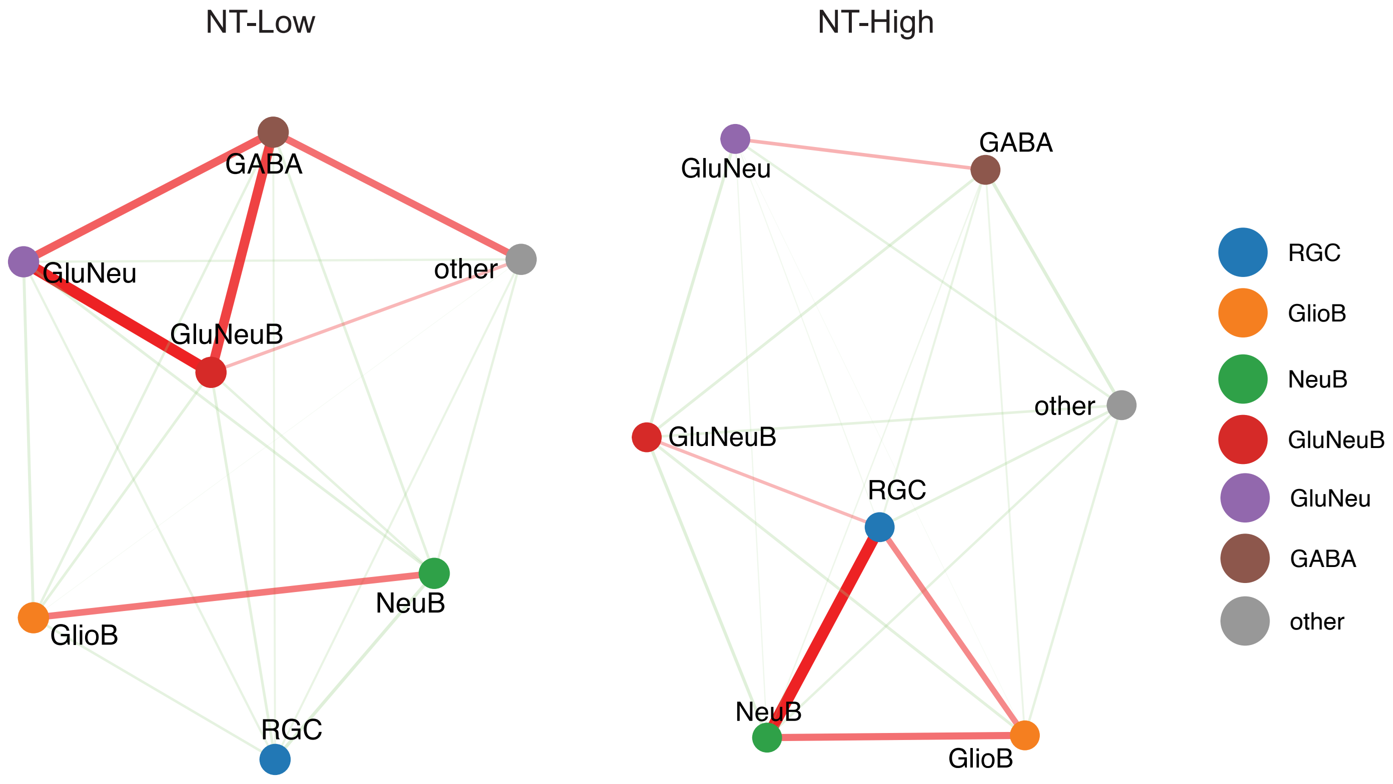
